# Joining Two Perspectives of the Proteome with the Trypsin+HTA Protease Workflow

**DOI:** 10.64898/2026.10.05.756883

**Authors:** Mario M. Alba, Ajay Bharadwaj, Lindsey Becker, Ethan J. Canfield, Jonathan E. Katz, Jennifer E. Van Eyk, Simion Kreimer

**Author notes:** **Corresponding Authors.** Jennifer E. Van Eyk, Smidt Heart Institute and Advanced Clinical Biosystems Research Institute, Cedars-Sinai Medical Center, Los Angeles, CA;. Simion Kreimer, Smidt Heart Institute and Advanced Clinical Biosystems Research Institute, Cedars-Sinai Medical Center, Los Angeles, CA.

## Abstract

Proteolysis can be completed in a single-step, rapid (5-30 minutes) reaction using hyperthermoacidic archaeal (HTA) proteases. Protocol simplicity and stability of HTA proteases in ambient storage conditions removes many logistical barriers to widespread deployment of proteomic analysis. HTA protease cleavage selectivity is complex but quantitatively reproducible and generates peptides that negligibly overlap with trypsin. Tryptic and HTA protease peptides emphasize different proteins and provide complementary coverage of amino acid sequence and post-translational modification (PTM) sites. Digestion of a sample’s separate aliquot with HTA protease is a minor addition to trypsin-based workflows, but LC-MS analysis of both sets of peptides is not practical for large cohorts. This manuscript evaluates the identifications and reproducibility of LC-MS analysis of tryptic and HTA protease peptide mixtures. Adding HTA protease peptides to 20% increased total protein identifications by 24% on average in neat plasma and identified 216 additional biologically interesting modifications compared to conventional trypsin-only bottom-up proteomic analysis. This approach is reproducible and does not increase sample preparation or analysis time. Peptides from each protease present independent opportunities to quantify overlapping proteins and a quantification model was developed in which, for each protein, the most complete protease data is selected for primary quantitation and data from the second protease is used to infer missing values. This cross-protease inference is more accurate than minimum, mean, or nearest neighbor imputations. Finally, biomarkers discovered through HTA protease peptides are amenable to rapid HTA protease only proteomic workflows thus streamlining translation from discovery to fieldable applications.

## Introduction

Hyperthermoacidic archaeal (HTA) proteases are derived from volcanic hot-spring dwelling single-cell Archaea and have optimal activity at pH 2-4 and 80-100 ºC; conditions that denature mesophile proteins^1^. Addition of a reducing reagent like 5 mM TCEP in the reaction buffer dissociates disulfide bonds. The cysteines remain protonated at the reaction pH and during acidic reversed phase chromatography, thus obviating a cysteine alkylation step. As such, raw input sample can be digested into peptides in one fast (5-30 min) reaction. As we^2^ and others^1,3^, previously reported proteolysis with the Krakatoa HTA protease manufactured by Cinder Biological Inc. is suitable for reproducible protein quantitation despite a complicated cleavage site selectivity. There is a unique population of proteins that are quantifiable with Krakatoa and not detected with trypsin proteolysis. This complementarity extends to protein amino acid sequence and post-translational modification sites (PTMs)^2^. Analyzing samples with each protease independently creates a more complete perspective, but it is expensive to double instrumentation usage, especially for large cohorts. In this manuscript we present and evaluate a workflow in which blood plasma samples are split, digested separately with trypsin and Krakatoa, and the peptides are mixed and analyzed together by liquid chromatography-mass spectrometry (LC-MS) (**Figure 1A**). There are multiple commercially available HTA proteases that optimally function in hot acidic conditions with non-tryptic cleavage selectivity that would also be suitable for this workflow, but this work exclusively utilized Krakatoa.

**Figure 1.**
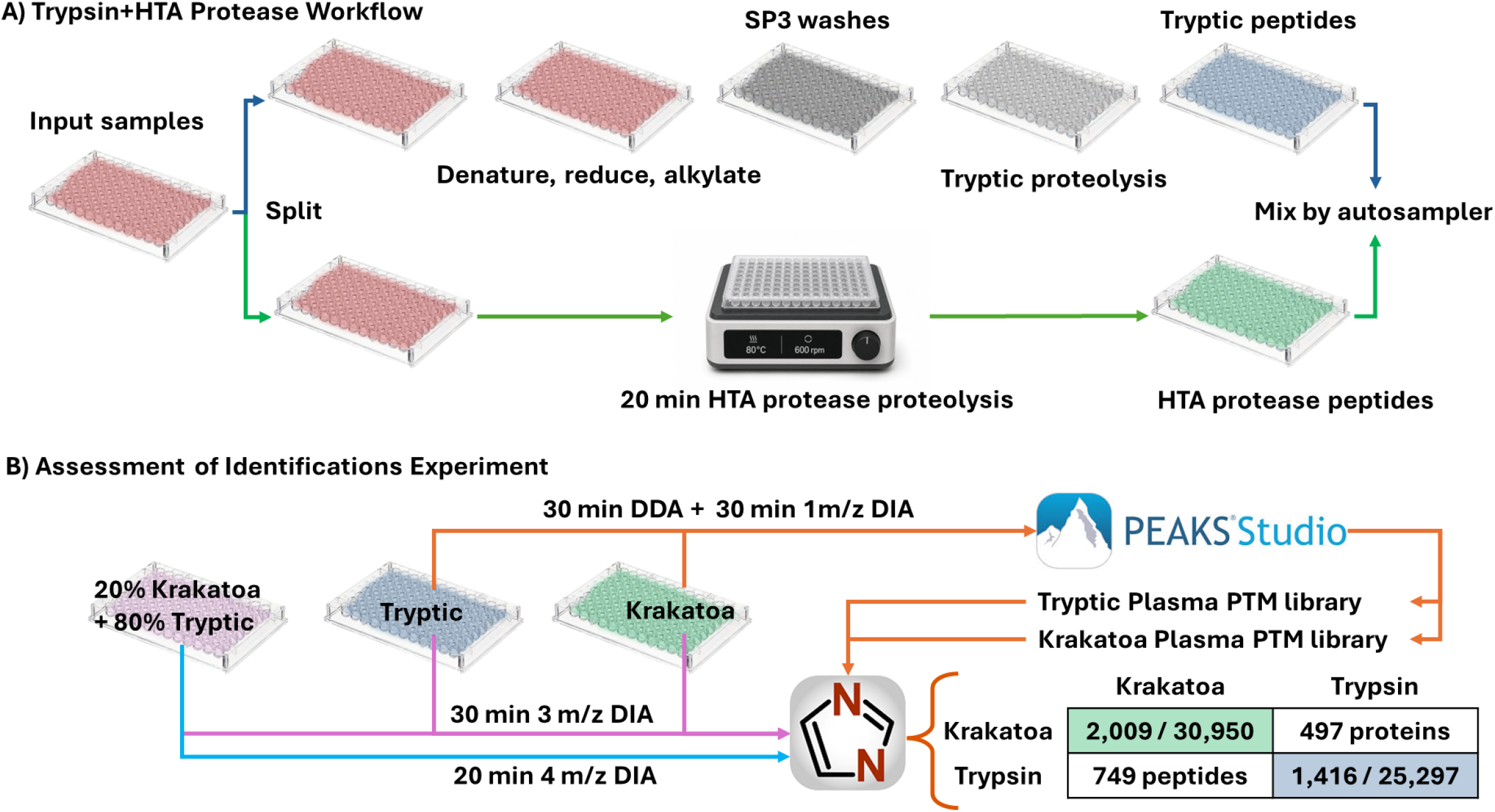
A – In the Trypsin+HTA Protease workflow the protein content of each sample is split to be digested separately with an automated SP3 protocol and a single-step HTA protease reaction. The peptides for each sample are then combined offline or on the autosampler before LC-MS analysis. B – Illustration of the Assessment of Identifications experiment in which the Krakatoa and Trypsin libraries were generated and used. Overlap of total identifications in the 54 samples from the two libraires (2,009 proteins and 30,950 peptides with HTA and 1,416 proteins, 25,297 with trypsin) was negligible for peptides (749) and only 497 for proteins.

Blood plasma is a noninvasively accessible source of protein biomarkers from all connected tissues and organs. Due to the overwhelming concentration of albumin and immunoglobulins the majority of plasma protein species are not detectable by LC-MS without enrichment^4,5^. The phenomenon of overly abundant species “masking” other analytes is attributed to limited mass analyzer dynamic range and ionization suppression^6,7^. Peptides have drastically varied response factors (signal intensities relative to concentration) due to differences in proteolysis recovery, chromatographic elution, ionization and fragmentation efficiency, and transmission to the mass analyzer^8^. Proteolysis with different cleavage specificities generates complementary sets of high responding peptides that increase the signals of proteins otherwise masked with a singular protease.

Trypsin is currently considered the default protease for bottom-up proteomics, but other proteases including chymotrypsin^9^, ProAlanase^10^, Thermolysin^11^, etc.^12^ with non-tryptic cleavage specificity can be used for quantitatively reproducible LC-MS analysis^11^. Peptides from these proteases added to conventional tryptic workflows would likely also increase the total quantifiable proteins. A related example is the use of multiple proteases in biotherapeutic characterization to achieve full sequence coverage^13^. Krakatoa proteolysis has two key advantages as an addition to tryptic sample preparations: (1) it can be completed quickly with minor effort and without increasing total sample preparation time and (2) there is negligible overlap in peptides with trypsin, which avoids peptide level signal convolution between the two proteases.

Commonly tryptic protocols include removal of process and sample contaminants, for example via solid phase extraction (SPE)^14^. For those protocols HTA protease peptides can be added before the clean-up to remove contaminants from both sets of peptides. In this manuscript, proteins were purified intact by precipitation with microparticles (SP3) before proteolysis^15^. Krakatoa peptides were added without removing plasma contaminants, which emulates a hypothetical field deployed Krakatoa-only workflow that would not involve offline sample clean-up. Instead, robust peptide trapping and elution chromatography was used to filter out hydrophilic contaminants and protect the analytical column from clog causing particulates.

Initial evaluation of workflow identifications used the trap and elute configuration with trapping column backflush on the Vanquish Neo nano liquid chromatography system (Thermo Fisher). The needle-seat clogged after continuous injection of Krakatoa and mixed peptide samples. Thus, a prototype LC system with a valve integrated dual trapping, single analytical column (DTSC) configuration created by ProteoPulse LLC. was used to evaluate workflow reproducibility. Two trapping columns allow a sample to be loaded and desalted on one trap during analysis of a sample from the alternate trap which eliminates data acquisition gaps between samples due to injection.^16,17^ Parallelizing sample analysis with subsequent sample loading also allows a sample to be manipulated for the duration of the analytical method without decreasing throughput. Krakatoa peptides were automatically mixed with tryptic peptides on the autosampler before each injection. Using the autosampler instead of manual peptide mixing allows for dynamic adjustment of the peptide ratio as desired. In this platform the trapping media was integrated directly into the 4-port valve rotor which achieved the same functionality as the 10-port valve from the first DTSC configuration with fewer potential points of failure. (**Figure S1**)

A major challenge of this workflow is interpretation of the data generated by data-dependent (DDA)^18^ and data independent acquisition (DIA)^19^ analysis. Identification and quantitation of tryptic peptides benefit from the mostly complete and specific cleavage of arginine and lysine residues. The complex cleavage specificity of the HTA protease currently demands the consideration of every residue as a potential cleavage site. Conventional false discovery rate (FDR) control generates a set of decoy sequences equal in size to the potential real peptide population and accepts candidate identifications above a score that includes a low percentage (*e*.*g*. <1%) of decoys relative to true peptide identifications^20^. When every residue is a potential cleavage site the decoy peptide population is enormous relative to the real peptide population in the sample and sporadic high scoring decoy matches reduce identification of HTA protease generated peptides. This effect is negated with the use of limited sample-specific peptide libraries in DIA data analysis^21,22^. However, this effect remains a limitation in creation of the peptide libraries. In this work, Human blood plasma libraries containing post-translational modifications (PTM) were generated for trypsin and Krakatoa peptides to identify similar numbers of proteins in both proteolysis conditions. (**Figure 1B**)

In this workflow peptides from each protease provides a channel of quantitative data for the overlapping proteins. We created a tool that joins the quantitative output matrices from DIA-NN data searches with each protease-specific peptide library. For each overlapping protein the algorithm selects the protease that quantified the protein in the most samples (lowest missingness). Then data from the alternate protease is scaled to infer missing protein quantities in the more complete protease data. This cross-protease inference is more accurate than conventionally used minimal^23^, mean^24^, or nearest-neighbor (kNN)^25^ value imputations. Thus, the Trypsin+HTA Protease workflow perceives more of the proteome by quantifying more protein species in more samples with increased amino acid sequence and PTM coverage than singular protease analysis.

## Materials and Methods

### Safety

Human blood plasma was handled under standard biosafety precautions without unexpected hazards.

### Ethical considerations

This study analyzed donated de-identified residual blood plasma with ethical approval from the Cedars-Sinai Medical Center IRB #STUDY0000169 Date Effective: 08/03/2021.

#### Tryptic Proteolysis

A 0.5 µL equivalent of dried blood plasma was reconstituted in 40 µL of 5% SDS, 10 mM TCEP, 40 mM chloroacetamide, 50 mM TEAB and incubated at 90 ºC for 10 min with sonication^15^. This achieved protein denaturation, and cysteine reduction and alkylation in a single step. The samples were cooled to room temperature and 40 µL of 8M urea in water were added to assist with removal of SDS. Acetonitrile was added to 70% and the rest of the protocol was automated on a Beckman i7 liquid handling system. 500 µg of ferrous core silica beads (G-Biosciences) were added in 5 µL of 50 mM Tris-HCl (pH 8) to precipitate the protein content onto the beads. The beads and bound proteins were magnetically immobilized while the samples were washed twice with 80% ethanol and twice with 100% acetonitrile. The beads were resuspended in 50 µL of 50 mM Tris-HCl at pH 8, 10 mM CaCl_2_ and 2 µg Trypsin/Lys-C (Promega). The samples were digested overnight at 37 ºC. The solvent was collected and a second elution using 50 µL of 4% acetonitrile 0.1% formic acid was combined with the first fraction.

#### Krakatoa Proteolysis

A 0.5 µL equivalent of blood plasma was resuspended in 50 µL of 5 mM TCEP, 60 mM phosphate citrate at pH 3 and digested with 10 units of Krakatoa for 20 minutes at 80 ºC. The second set of Krakatoa peptides from the Assessment of Identifications experiment was also incubated with 40 mM chloroacetamide for 30 min so that carbamidomethylated Krakatoa peptides could be included in the generated library. The first Krakatoa sample preparation did not include cysteine alkylation.

#### Assessment of Identifications

Aliquots of 0.5 µL from a random set of 54 residual blood plasma samples were prepared twice with Krakatoa and twice with trypsin. Peptides from the second pair of proteolysis reactions were mixed at 20% Krakatoa to 80% trypsin ratio to generate the mixed samples. The samples were analyzed on an Orbitrap Astral MS and Vanquish Neo LC (Thermo) configured for trap-and-elute with trapping column backflush.

Peptide libraries were created from 30 min (**Figure S2A**) DDA analysis of the first set of preparations and 1 m/z isolation window DIA analysis (**Figure S3**) of the second set of preparations. All samples were analyzed using the quantitative 30 min, 3 m/z isolation window DIA method with the same liquid chromatography separation. The peptide mixtures were analyzed with both the 30 min method and a 20 min, 4 m/z isolation window DIA method (**Figure S2B**).

#### Assessment of Reproducibility

A set of 8 de-identified residual plasma samples were digested in 8 replicates on 3 different days (24 preparations per sample) using the tryptic SP3 protocol and Krakatoa on separate 96-well plates. The plates were placed into the prototype autosampler from ProteoPulse LLC and mixed on-line in equal portion before injection. The combined peptides were separated using a 15 min method (**Figure S2C**) and analyzed using 30 m/z wide isolation window DIA on a Orbitrap Exploris 480 (**Figure S3**). The data were processed in DIA-NN 2.3.1 using the libraries generated in the Assessment of Identifications experiment.

## Data Analysis

DDA and 1 m/z isolation window DIA data were analyzed separately in PEAKS 13.1 using the DDA workflow^26,27^. PEAKS identifies the most probable precursor masses from the isolation window, which allows processing of 1 m/z isolation window DIA data as if it were acquired by DDA. The precursor and fragment mass error tolerances were set to 6 ppm and 0.01 Da, respectively. False discovery rates (FDR) were controlled at 1% at the peptide spectral match (PSM) and protein group levels. All built-in modifications were considered for PTM analysis with *de novo* ALC threshold set to 15%. Peptide lengths were set to 6-60 residues. The tryptic data were analyzed with semi-tryptic specificity and two maximum missed cleavages. The Krakatoa data were analyzed with no protease specificity. Complete carbamidomethylation of all cysteines was set for the tryptic data and the 1 m/z Krakatoa data (samples were alkylated after the standard preparation), but not the DDA Krakatoa data (samples were not alkylated).

Four libraries were compiled in PEAKS: Krakatoa 1 m/z DIA, Krakatoa DDA, Tryptic 1 m/z DIA, and Tryptic DDA. The libraries were formatted and merged for each protease using in-house developed R-scripts. The DIA data were analyzed in DIA-NN separately with each library (**Figure 1B**)^28^. For each protein quantified in both the Krakatoa and trypsin library searches the cross-protease inference algorithm selected the search in which the protein was quantified in the greater fraction of samples. A Kendall’s τ test was then used to confirm that the two proteases’ measurements were correlated across samples for the protein and that missing measurements in one search could be inferred from the second search. In samples where the elected protease was missing a value, the values was inferred by scaling the second protease’s measurement of that protein in the same sample by the log transformed median difference between the two proteases for that protein across all samples (**Figure S4**).

## Results and Discussion

The generated Krakatoa and trypsin protease-specific peptide libraries were implemented in DIA data analysis and showed negligible overlap (749) between trypsin (25,297) and Krakatoa (30,950) peptides. This complementarity extended to protein identifications with an overlap of 497 between the 2,009 Krakatoa and 1,416 trypsin proteins (**Figure 1B**). The addition of 20% Krakatoa to 80% trypsin peptides increased the protein identifications by 24% on average (**Figure 2A** Trypsin 2 vs. Mix 30). The number of identifications matched between repeated sample preparations with high overlap (Krakatoa 1 vs. 2 and Trypsin 1 vs. 2). A similar number of proteins were detected with the 20 min method (Mix 20) as with the 30 min trypsin-only analysis (**Figure 2A**). The increased dynamic range of narrow window DIA on the Orbitrap Astral resulted in comparable numbers of identified proteins between Krakatoa and Trypsin, whereas in our previous work on an Exploris 480 (Thermo Fisher) and timsTOF Pro 2 (Bruker), Krakatoa identified half as many proteins as trypsin^2^.

**Figure 2.**
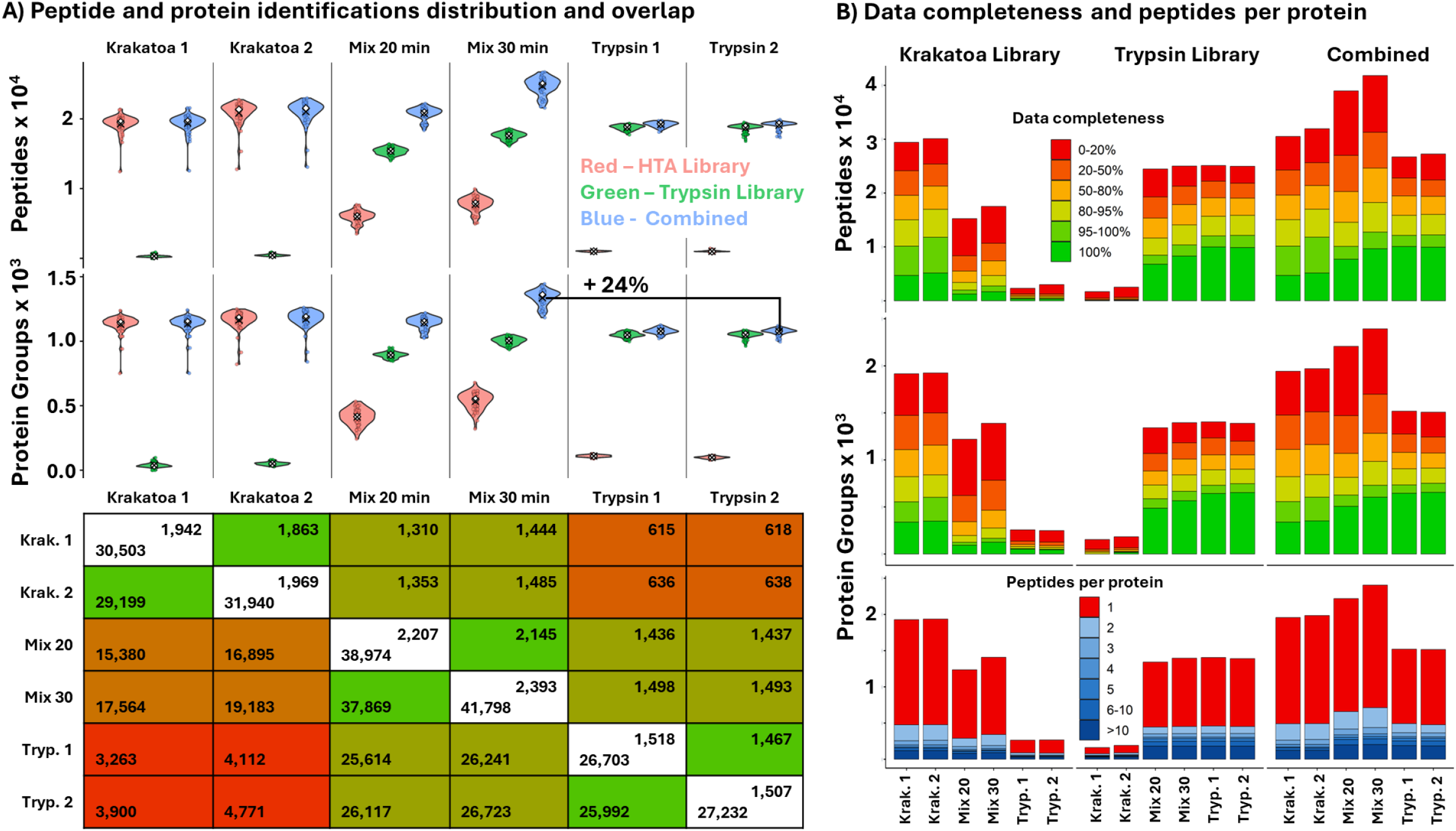
A – The identifications within the samples (n=54) using the tryptic and Krakatoa peptide libraries (top). Identifications between repeated Krakatoa and Trypsin preparations (1 and 2) was reproducible. Predictably, few proteins and peptides were identified when tryptic samples were analyzed against the Krakatoa library and *vice versa*. When 20% Krakatoa peptides are mixed with tryptic peptides (Mix 20 min and Mix 30 min) 262, or 24% more proteins are identified on average than the Trypsin 2 identifications alone. Notably a 20-minute analysis of the mixture identified the same number of proteins species as the 30-minute analysis of the Trypsin 2 samples. The matrix shows the identification overlap between sample preparations, protein on the top right and peptide on the left bottom. The diagonal shows the total identifications within each preparation and overlap is represented by color (green – high to red – low). B – The completeness (% of samples where the peptide (top) or protein (middle) are identified. The bottom plot presents the number of peptides identified per protein.

The ratio of 20% Krakatoa to 80% trypsin peptides based on input material was used to minimally perturb the tryptic data while increasing the total number of identified proteins. Of the 1,507 proteins and 27,232 peptides identified in the second tryptic set of peptides 1,493 and 26,723 were detected in the mixture. (**Figure 2A**) Quantitation was also minimally perturbed and the correlation (R^2^) between the mixture and the second trypsin set (Mix 30 vs. Trypsin 2) was slightly higher than repeated trypsin proteolysis (Trypsin 1 vs. Trypsin 2) (**Figure S5**). Thus, the addition of Krakatoa peptides at 20% had a negligible impact on the trypsin-based perspective of the proteome.

Along with quantifying more protein species, the Krakatoa peptides also increased the amino acid sequence and modification site coverage of many proteins. Of the 1,493 overlapping proteins, sequence coverage increased for 368 proteins, and was reduced for 12. Of the 1,262 biologically interesting modifications detected in the second trypsin set, 1,242 were still detected in the peptide mixture, and 236 Krakatoa modifications were added. Most of these modifications were also detected with the 20 min method (**Figure 3A**). **Figure 3B** illustrates how the amino acid coverage and modification information from the two proteases is combined in the peptide mixture analysis for three representative proteins. Notably, the number of modifications identified by trypsin in the generated libraries is three times higher than Krakatoa. This pattern also holds for modifications that are possible artifacts of sample processing (**Figure S6**). It is unlikely that proteolysis with Krakatoa uniformly removes 2/3 of all modifications. We attribute this difference to reduced sensitivity in PTM identification from DDA data when the peptide libraries are generated. Development of data processing algorithms tailored to HTA proteases should find more modifications in this data and future experiments.

**Figure 3.**
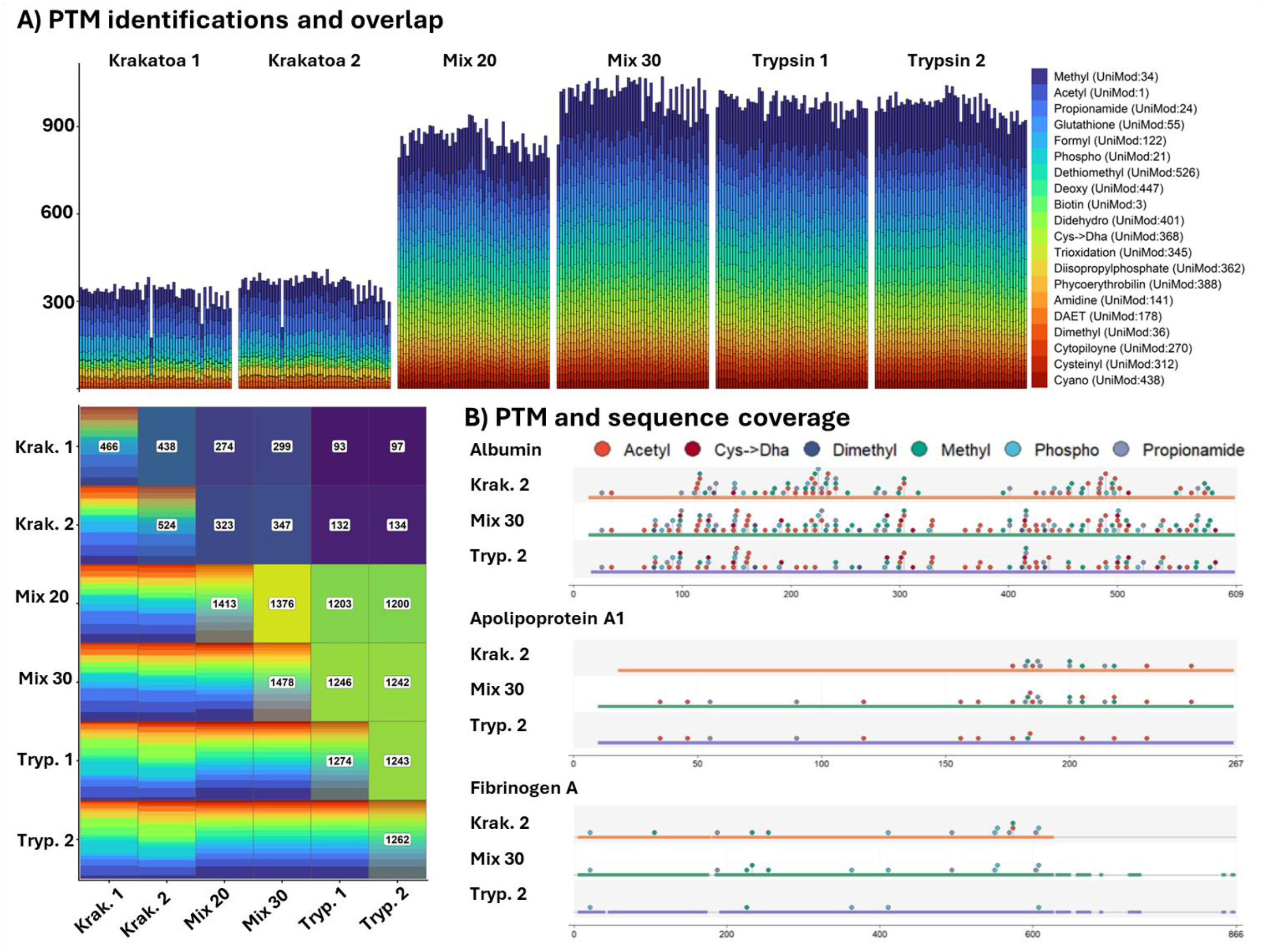
A – PTM sites quantified in each sample. Each stacked bar represents the number of specific modifications, indicated by color, in each sample by preparation group. The matrix shows the overlap in PTM sites between preparation groups with specific modification indicated by color. B – Maps of sequence coverage and detected modifications for three proteins, each dot represents a specific modification on a given site, the bar represents the amino acid sequence covered by the peptides generated by each protease (X-axis indicates the residue position in the sequence). Mix 30 combines peptides from Krakatoa 2 and Trypsin 2, which increases coverage and detectable PTMs.

The quantitative reproducibility of the workflow was tested using an equal mixture of peptides by repeated processing of 8 aliquots from 8 plasma samples for 3 nonconsecutive days. The equal peptide ratio was selected to evenly compare the reproducibility of Krakatoa and tryptic sample processing. The measured variance combined the variability of instrument performance, autosampler sample mixing, and loading of the mixed samples onto the valve rotor integrated trapping columns. A throughput of 96 samples per day (SPD) was selected to match the pace of one 96-well plate prepared per day. Due to the faster method and the lower dynamic range of the Exploris 480 DIA compared to the narrow window Orbitrap Astral DIA fewer proteins and peptides were identified in this experiment. The proteins quantified in all 24 replicates of a sample with an inter-day CV <20% were considered as well-quantified. This number was sample dependent, and 38-60 proteins were well-quantified with Krakatoa peptides while 56-107 were quantified with tryptic peptides. The majority of these proteins were highly abundant species represented in both peptide sets, nonetheless the combined number of well quantified proteins was 73-119 before cross-protease quantitation (**Figure 4A**).

**Figure 4.**
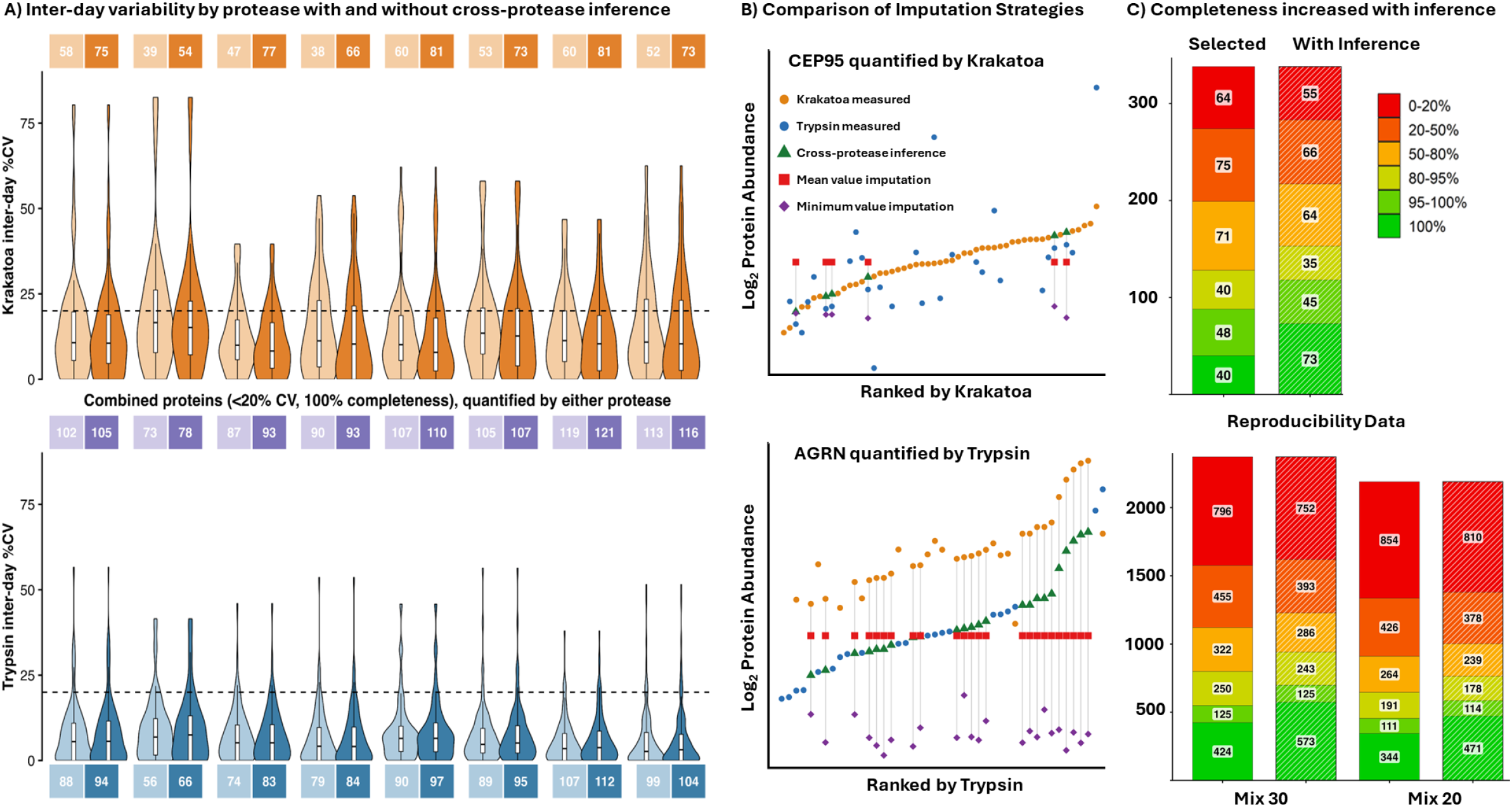
A – Coefficient of variance distributions (violin plots) over 8 replicate preparations across three days measured with Krakatoa (orange) and trypsin (blue). The box number indicates the number of proteins quantified in all 24 samples with a CV <20%. The purple boxes represent the total well-quantified proteins from both proteases. The darker shading is with cross-protease inference enabled. B – Visualization of cross-protease inference and mean and minimum value imputations for two proteins. C – Impact of cross-protease inference on data completeness from the two datasets. The stacked bars show the completeness of quantitation using the protease with the most complete data, the data with stripes shows completeness with datapoints recovered using cross-protease inference.

**Figure 4B** illustrates quantitation for two proteins across the 54 samples in the Assessment of Identifications data sets. In the top panel CEP95 is more completely quantified with Krakatoa peptides (orange circles), but there are 6 samples in which the protein was not quantified with Krakatoa. The bottom panel shows that AGRN is more completely quantified with trypsin (blue circles) even though AGRN was not quantified by trypsin in almost half of the samples. The missing data could be imputed with a sample minimum value (purple diamond) or a dataset mean value (red square). But these strategies disregard the information from peptides generated by the alternate protease. Quantified peptides from the alternate protease indicate that the protein is present at detectable quantities, which invalidates a minimal value imputation. Differences in the alternate peptide quantities indicates that a single dataset mean value imputation is not accurate for every sample.

Data point recovery strategy accuracies were compared by masking 30% of measured values in the Assessment of Identifications dataset, replacing the missing values using each strategy, and then evaluating the mean error between the masked and filled values. In this test the cross-protease inference outperformed the imputation strategies (**Figure S7**). When applied to the reproducibility data, cross-protease inference increased data completeness to 100% for 2-6 proteins while maintaining the CV <20% thus increasing the number of well-quantified species (**Figure 4A**). In the Assessment of Identifications data cross protease inference increase the number of proteins quantified in all samples from 424 to 573 in the Mix 30 data and from 344 to 471 in the Mix 20 data (**Figure 4C**). It’s important to note that **Figure 4C** represents protein quantifications and thus differs from **Figure 1B** which represents protein identifications. With cross-protease inference missing protein quantities can be recovered accurately and a more complete perspective of proteomes across samples is achieved.

## Conclusion

The Trypsin+HTA Protease workflow joins the complementary perspectives of the proteome afforded by the two sample preparations. Trypsin benefits from reproducible and mostly complete cleavage of lysine and arginine residues and has nearly monopolized bottom-up proteomics for decades. This singular perspective has overlooked proteins and regions of proteins with too many, too few, or modification obscured cleavage sites. In proteomes with a high concentration range, like blood plasma, limited instrumentation dynamic range biases the detectable proteome to high concentration proteins and low concentration proteins that consistently proteolyze into high responding tryptic peptides. HTA Proteases have a complex cleavage specificity that is fully complementary to trypsin thereby exposing novel proteins, protein regions, and modifications. HTA Proteases have the added benefit of simple, rapid, and fieldable sample preparation, providing an easy addition to conventional sample preparation workflows. Mixing the peptides from tryptic and HTA Protease workflows joins the two perspectives of the proteome to provide a more complete view by increasing total quantified protein species and post-translational modifications, and accurately recovering missing protein quantities using cross-protease inference. Widespread implementation of the Trypsin+HTA Protease workflow, will discover biomarkers quantifiable with HTA Protease peptides which will benefit from easy translation into rapid fieldable HTA Protease only assays.

## Supporting Information

Figure S1. Valve diagrams of original DTSC and valve integrated DTSC, Figure S2. LC method details, Figure S3. MS method details, Figure S4. Cross-protease imputation schematic, Figure S5. Quantitative correlation between trypsin preparations and peptide mixture, Figure S6. Distribution of modifications that are likely artifacts of sample preparation, Figure S7. Masking test comparing minimum, median, and cross-protease imputation

## Data Availability

All collected data are available through the Massive repository (ID: MSV000103457). Reviewer user name MSV000103457_reviewer, password: Massive@JVE2026. The developed scripts and peptide libraries are shared through Github at https://github.com/ProteoPulse/Two-Protease-Paper

## Author Contributions

M.M.A. prepared the samples, analyzed the data, and assisted with the manuscript. A.B. assisted with analysis of the data and development of the cross-protease imputation strategy. A.B and M.M.A. are first co-authors. S.K. developed the methodology and instrumentation, conceived and designed the study and wrote the manuscript. L.B. assisted with sample preparations. E.J.C. and J.E.K. provided the samples. J.E.V.E. supervised the work and provided feedback and funding. S.C.L. provided mentorship for M.M.A. and funding. J.E.V.E. and S.K. are corresponding authors. All authors reviewed and approved the manuscript.

## Acknowledgments

We thank Steven Yannone at Cinder Biological Inc. for helpful discussion and providing the Krakatoa protease partially funded through National Institute of General Medical Sciences of the National Institutes of Health under award number 5R44GM128540. This work was mainly supported by funding to J.E.V.E. NIH #5U01DK127403-03, #5U01DK124019-04, and 1R01HL180612-01A1, and the Post-Doctoral Fellowship grant for Mario Alba (T32HL170963-03). Additional thanks to the Cedars-Sinai Proteomics and Metabolomics Core for instrument access. J.E.V.E. is the Erika Glazer Endowed Chair in Women’s Heart Health.

