## Supplementary material for "Joining Two Perspectives of the Proteome with the Trypsin+HTA Protease Workflow": Supplemetal Information

**Table of Contents**

Figure S1. Valve diagrams of original DTSC and valve integrated DTSC	S2

Figure S2. LC method details	S3

Figure S3. MS method details	S4

Figure S4. Cross-protease imputation schematic	S5

Figure S5. Quantitative correlation between trypsin preparations and peptide mixture	S6

Figure S6. Distribution of modifications that are likely artifacts of sample preparation	S7

Figure S7. Masking test comparing minimum, median, and cross-protease imputation	S8

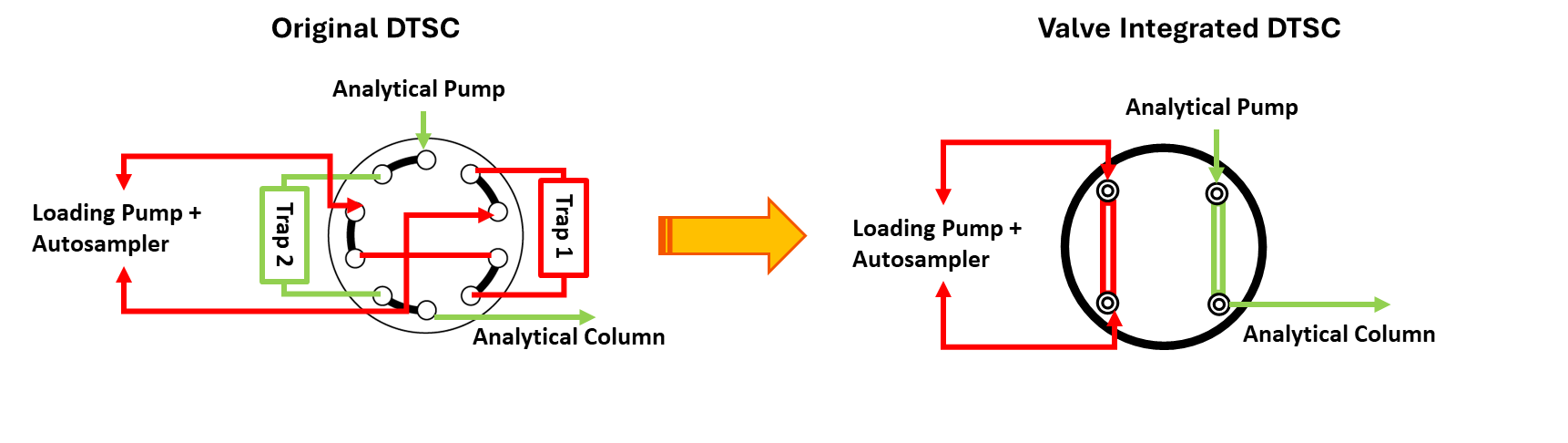

**Figure S1.** Valve diagrams of original dual-trap single column (DTSC) and valve integrated DTSC. DTSC allows for one trapping column to be flushed and equilibrated in both direction loaded with the subsequent sample and desalted all while the previous sample is running. The valve orientation is switched at the beginning of the run to start the next sample which eliminates gaps in data acquisition due to sample injection. Integrating the trapping media directly into the valve rotor instead of using external trapping columns substantially reduces the number of connections and potential points of failure while retaining the same functionality.

| **A) 30-minute Vanquish Neo method** | | |  | **B) 20-minute Vanquish Neo method** | | |  | **C) 15-minute ProteoPulse method** | | |
| --- | --- | --- | --- | --- | --- | --- | --- | --- | --- | --- |
| **Time (min)** | **Flow (nL/min)** | **%B** |  | **Time (min)** | **Flow (nL/min)** | **%B** |  | **Time (min)** | **Flow (nL/min)** | **%B** |
| 0 | 1200 | 4 |  | 0 | 1200 | 4 |  | 0 | 2500 | 12 |
| 2 | 1200 | 9 |  | 2 | 1200 | 9 |  | 1.2 | 2500 | 15 |
| 18 | 1200 | 25 |  | 13 | 1200 | 25 |  | 11.7 | 2500 | 36 |
| 27 | 1200 | 35 |  | 16 | 1200 | 35 |  | 12.7 | 2500 | 65 |
| 27.4 | 3000 | 99 |  | 17.5 | 1200 | 50 |  | 12.9 | 4000 | 99 |
| 27.9 | 3000 | 99 |  | 17.7 | 3000 | 99 |  | 13.3 | 4000 | 99 |
| 28.1 | 3000 | 5 |  | 18.2 | 3000 | 99 |  | 13.4 | 4000 | 12 |
| 28.3 | 3000 | 99 |  | 18.4 | 3000 | 5 |  | 13.5 | 4000 | 99 |
| 28.8 | 3000 | 99 |  | 18.5 | 3000 | 99 |  | 14 | 4000 | 99 |
| 29 | 3000 | 5 |  | 19 | 3000 | 99 |  | 14.1 | 4000 | 12 |
| 29.2 | 3000 | 99 |  | 19.2 | 3000 | 5 |  | 14.3 | 4000 | 12 |
| 29.7 | 3000 | 99 |  | 19.3 | 3000 | 99 |  | 14.5 | 2500 | 12 |
| 29.9 | 1000 | 5 |  | 19.8 | 3000 | 99 |  | 14.7 | 2500 | 12 |
|  |  |  |  | 19.9 | 1000 | 5 |  |  |  |  |
| **Trapping column:** Exp 2 cartridge trapping column 170 nL volume packed with PLRP solid phase (Optimize Technologies) | | | | | | |  | **Trapping columns:** Valve Integrated | | |
| **Analytical column:** 150 µm internal diameter, 15 cm length, 1.5 µm C18 particles PepSep (Bruker) | | | | | | |  | **Analytical column:** 150 µm internal diameter, 8 cm length, 1.5 µm C18 particles PepSep (Bruker) | | |

**Figure S2.** Details of the liquid chromatography methods used in the manuscript. %B is the proportion of organic buffer B (80% acetonitrile, 20% water 0.1% formic acid) in aquaeous buffer (100% water, 0.1% formic acid). The 30 minute and 20 minute methods used the same components.

|  | **3 m/z DIA** | **4 m/z DIA** | **1 m/z DIA** | **DDA** | **Reproducibility** |
| --- | --- | --- | --- | --- | --- |
| Instrument | Orbitrap Astral | Orbitrap Astral | Orbitrap Astral | Orbitrap Astral | Exploris 480 |
| Acquisition | DIA | DIA | DIA | DDA | DIA |
| Method duration (min) | 30 | 20 | 30 | 30 | 15 |
| Spray voltage (V) | 2300 | 2500 | 2300 | 2300 | 2900 |
| Ion transfer tube (°C) | 280 | 280 | 280 | 280 | 280 |
| RF lens (%) | 40 | 50 | 50 | 50 | 50 |
|  | **MS1 Settings** | | | | |
| MS1 resolution | 240,000 | 240,000 | 240,000 | 240,000 | 60,000 |
| MS1 range (m/z) | 360–1200 | 300–1000 | 300–1200 | 300–1400 | 380–1200 |
|  | **MS 2 Settings** | | | | |
| MS2 isolation window (m/z) | 3 | 4 | 1 | 1.2 | 30 |
| Number of scan events | 200 | 150 | 600 | top-50 | 17 |
| Precursor range (m/z) | 380–980 | 380–980 | 380–980 | 300-1400 | 400–900 |
| MS2 resolution | Astral | Astral | Astral | Astral | 15,000 |
| MS2 max injection time (ms) | 2.7 | 4 | 2.7 | 5 | 25 |
| MS2 normalized AGC (%) | 200 | 200 | 200 | 200 | 400 |
| HCD collision energy (%) | 30 | 30 | 30 | 30 | 30 |
| Cycle time | 0.4 s | 0.5 s | 0.4 s | top 50 | 3 s |

**Figure S3.** Details of the mass spectrometry methods used in the manuscript.

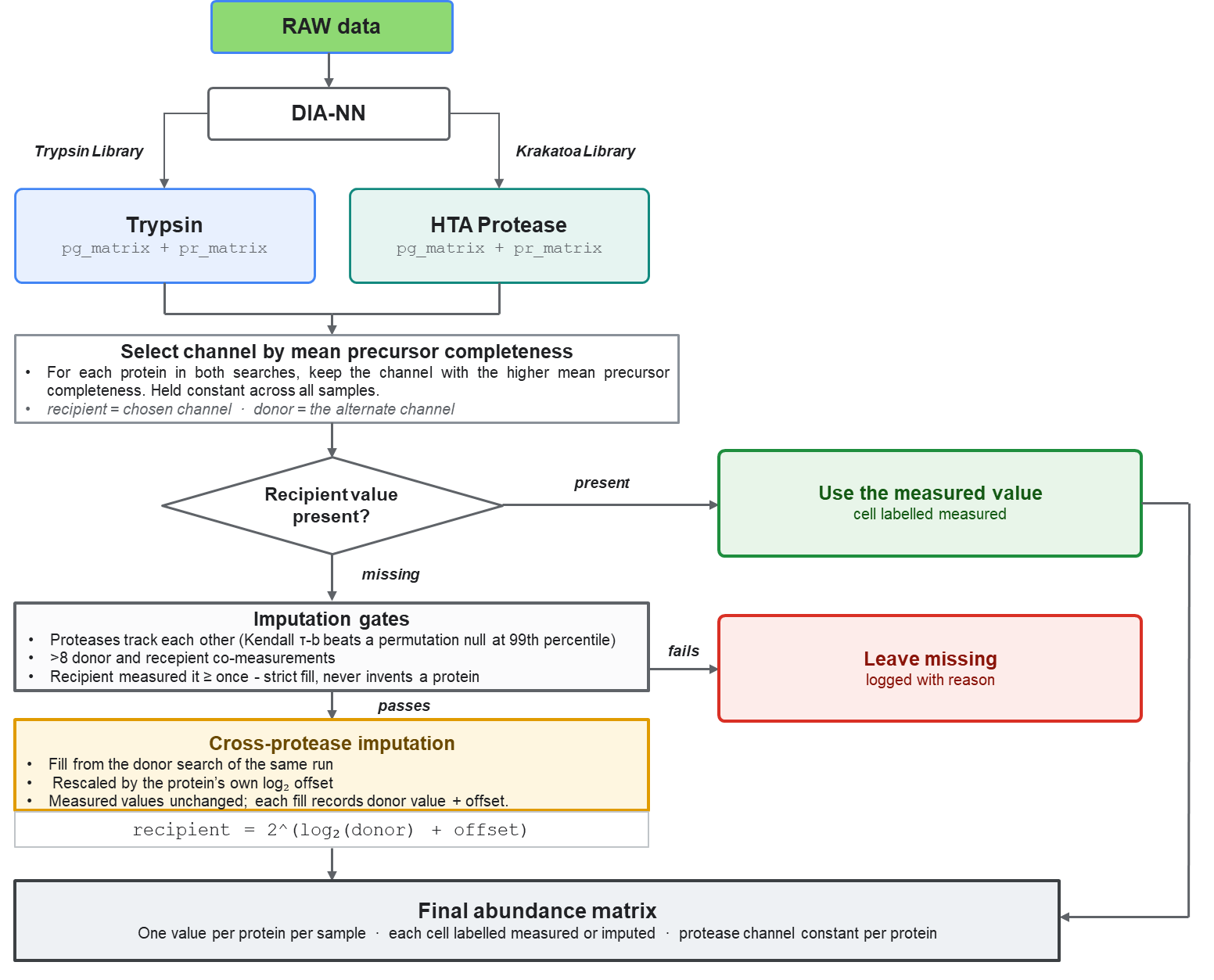

**Figure S4.** Schematic of the cross-protease data imputation strategy in QuantAdvisor. Starting with the DIA-NN output from analyzing the raw data separately with each of the libraries QuantAdvisor finds the proteins quantified in both sets of results and selects the protease that most completely quantifies each protein across the samples. If there are missing values in the selected or recepient data three tests determine if cross-protease imputation should occur. The key test is a Kendal’s Tau score that suggests that the two proteases generally agree on quantifying the protein in question. If the conditions are met , cross protease quantitation occurs by scaling the donor value using an offset in the logarithmic space (recepient = 2 ^ (log_2_ (donor) + offset). The imputed and measured values are then compiled into a final abundance matrix, with one set of quantities for each protein.

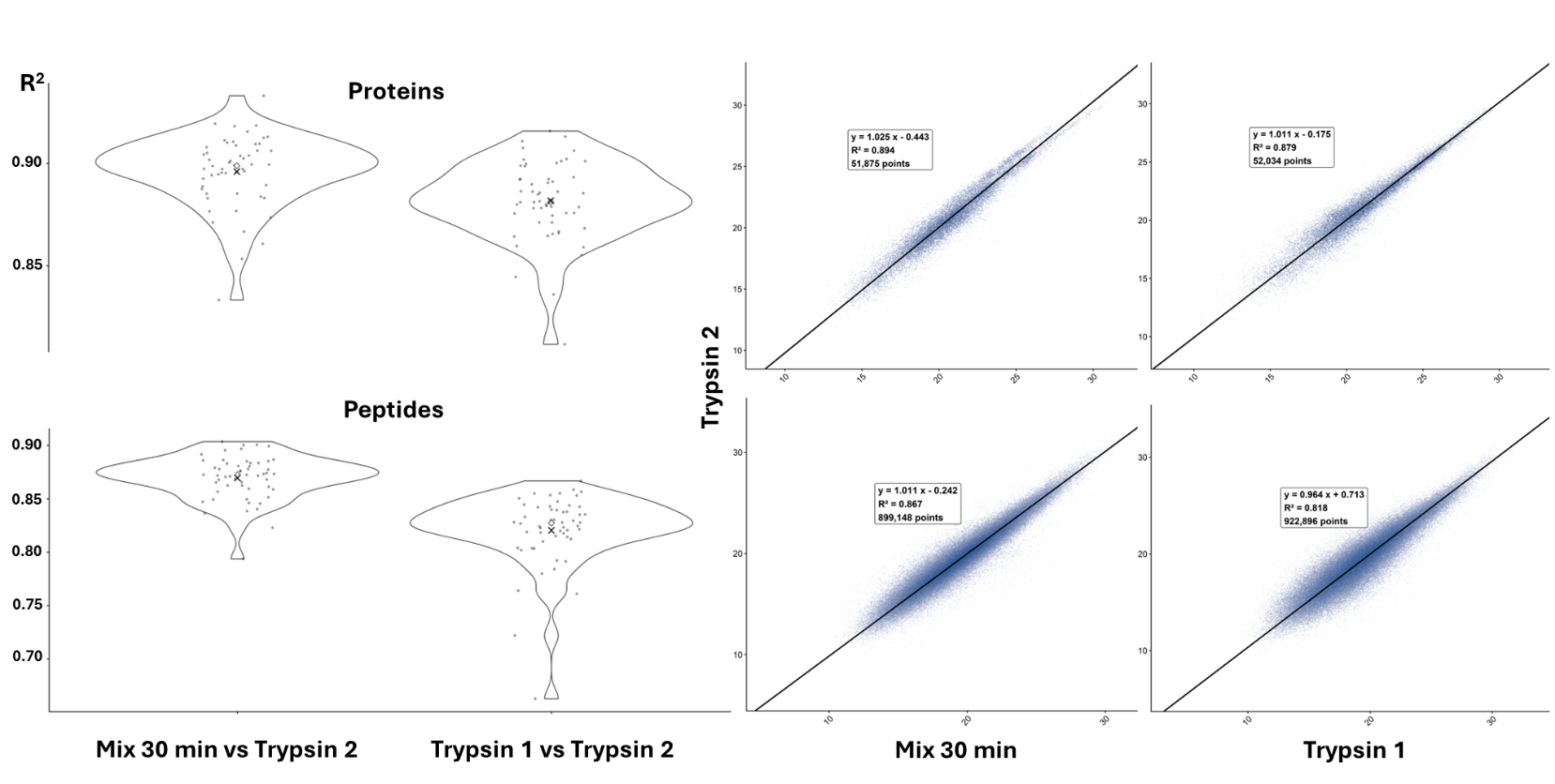

**Figure S5.** Quantitative correlation between a 20% Krakatoa 80% trypsin peptide mixture and 100% trypsin. The violin plots represent the R^2^ values for each pair of samples at the protein (top) and peptide (bottom) level. The scatter plots show the correlation of all datapoints pairs across all samples and their correlation. The correlation between the peptide mixture and 100% trypsin peptides (Mix 30 vs. Trypsin 2) is higher than two separate prepaprations (Trypsin 1 vs. 2).

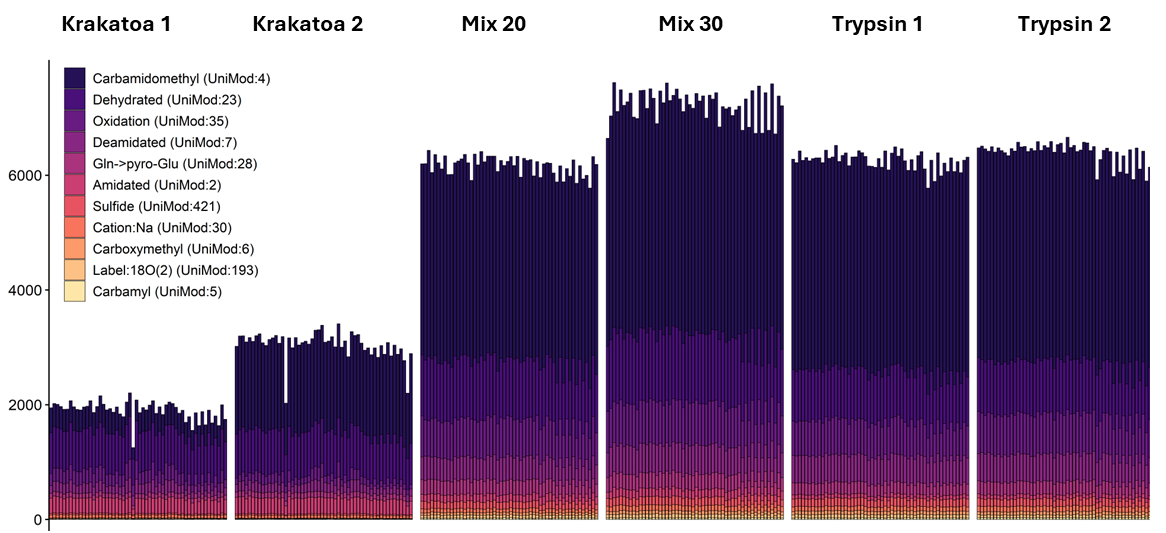

**Figure S6.** Peptide modifcations identified in each sample by sample preparation strategy that are possible artifacts of sample handling. As with the biologically interesting PTMs roughly 3 times as many modifications are identified in the trypsin preparations as the Krakatoa preparations, suggesting the reduced number is due to data processing limitations. The second Krakatoa preparation included alkylations, reflexting the large number of carbamidomethylated peptides.

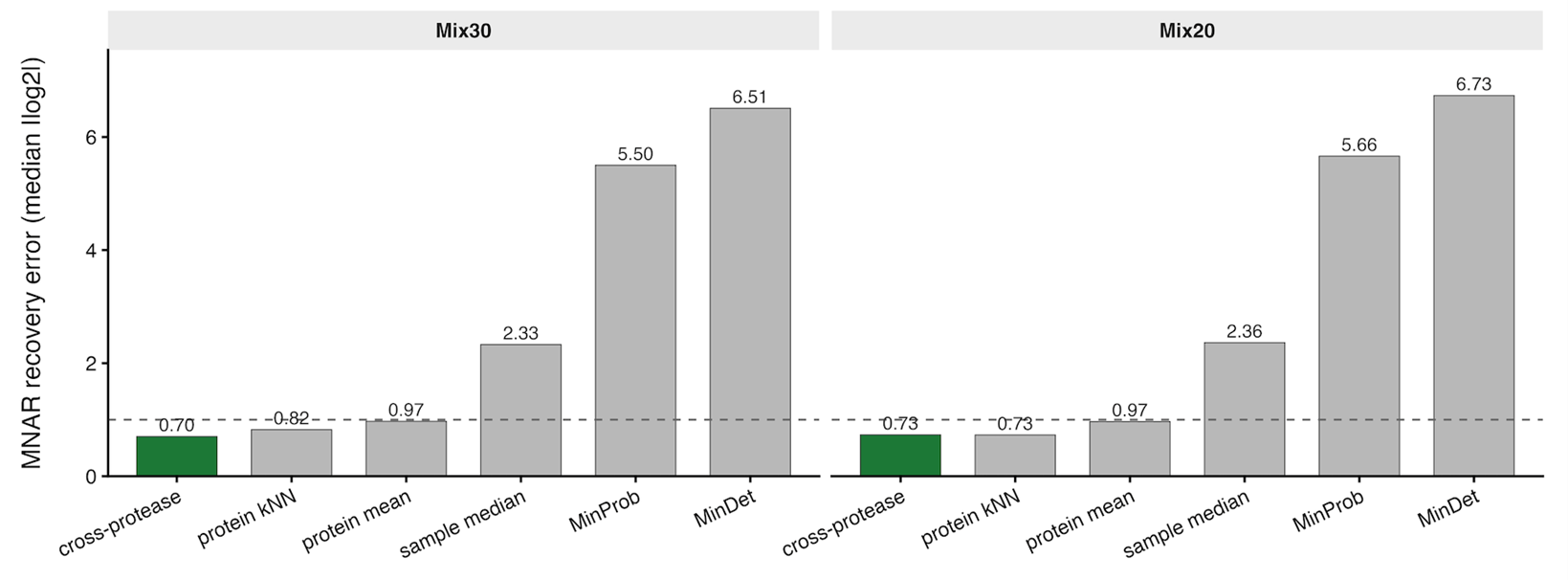

**Figure S7.** Results of the masking experiment to compare the accuracy of imputation strategies. In this experiment 30% of protein quantities in the Mix 30 and Mix 20 results from the Assessment of Identifications experiment were masked and replaced using each imputation strategy the mean error between the masked value and the replaced value is used as a gauge of accuracy and a lower error is better. Cross-protease inference is more accurate than common imputation strategies.
